# Modulation of bacterial cell size and growth rate via activation of a cell envelope stress response

**DOI:** 10.1101/2022.07.26.501648

**Authors:** Amanda Miguel, Matylda Zietek, Handuo Shi, Anna Sueki, Lisa Maier, Jolanda Verheul, Tanneke den Blaauwen, David Van Valen, Athanasios Typas, Kerwyn Casey Huang

## Abstract

Fluctuating conditions and diverse stresses are typical in natural environments. In response, cells mount complex responses across multiple scales, including adjusting their shape to withstand stress. In enterobacteria, the Rcs phosphorelay is activated by cell envelope damage and by changes to periplasmic dimensions and cell width. Here, we investigated the physiological and morphological consequences of Rcs activation in *Escherichia coli* in the absence of stresses, using an inducible version of RcsF that mislocalizes to the inner membrane, RcsF^IM^. Expression of RcsF^IM^ immediately reduced cellular growth rate and the added length per cell cycle in a manner that was directly dependent on induction levels, but independent of Rcs-induced capsule production. At the same time, cells increased intracellular concentration of the cell division protein FtsZ, and decreased the distance between division rings in filamentous cells. Depletion of the Rcs negative regulator IgaA phenocopied RcsF^IM^ induction, indicating that IgaA is essential due to growth inhibition in its absence. However, A22 treatment did not affect growth rate or FtsZ intracellular concentration, despite activating the Rcs system. These findings suggest that the effect of Rcs activation on FtsZ levels is mediated indirectly through growth-rate changes, and highlight feedbacks among the Rcs stress response, growth dynamics, and cell-size control.

## Introduction

The environment plays a key role in determining the physiological and physical state of a bacterial cell. In natural conditions, bacteria are exposed to varied and often stressful environments that can include changes to pH, temperature, and nutrient availability, as well as the presence of harmful chemicals such as antibiotics. The myriad ways that bacteria respond to such environmental changes include modulation of growth rate, protein composition, cell envelope modification, and cell size and shape [1, 2]. Many of these changes in behavior are governed by complex signaling pathways, each with their own set of triggers. These triggers can overlap; for example, as *Escherichia coli* cells encounter starvation conditions, they express the master transcriptional regulator RpoS, which activates programs involved in osmotic, oxidative, and envelope stress [3, 4]. Due to the overlap of these stress responses, it remains an open question as to how many response pathways specifically alter cellular physiology.

The Rcs phosphorelay is a stress response pathway that responds to damage in the cell envelope of Gram-negative bacteria [5, 6]. In *E. coli*, deletion of genes in the Rcs pathway causes sensitivity to cell shape-perturbing drugs such as A22 and mecillinam [7], suggesting that the Rcs system may play a yet unidentified role helping cells adjust to shape changes. The Rcs system senses envelope stress via the outer membrane lipoprotein RcsF. Under normal growth conditions, RcsF is transported to the outer membrane and ultimately surface exposed [8, 9].

However, when its transport to the outer membrane or surface exposure is perturbed due to envelope stress, RcsF remains in the periplasm, free to interact with the inner membrane protein IgaA [8, 10, 11], and thereby to activate the Rcs system [12]. When activated, the Rcs system regulates the expression of genes controlling many functions, including production of the exopolysaccharide colanic acid [13]. A previous study suggested that the Rcs system transcriptionally regulates cell division proteins, further supporting a connection between Rcs activation and cell shape [14]. A mutant variant of RcsF that localizes to the inner membrane (RcsF^IM^) constitutively activates the Rcs pathway [8, 15], providing a straightforward mechanism for induction of the Rcs system in the absence of environmental stress.

Environmental changes can have varied impacts on bacterial cell shape and size, features that are connected with behaviors such as motility, adhesion, and immune evasion [16], all of which are regulated by the Rcs pathway [6]. The nutritional content of the environment also strongly affects *E. coli* and *Salmonella enterica* cell size, with faster growing cells having larger volume [2, 17]. In transitions between nutrient-poor and nutrient-rich environments, *E. coli* cells rapidly adjust their length and width within an hour [18, 19]. By contrast, during steady-state exponential growth, bacteria robustly maintain their cell shape and size [20]. This cell-size homeostasis results from an adder mechanism of growth [21, 22], in which cells on average add a fixed volume Δ*V* during each cell cycle [23]. However, the molecular mechanisms by which cells modulate their cell size during nutrient changes and how they are coupled to growth rate remain largely unknown.

Bacterial cell size and shape is determined by the cell wall, a single macromolecule composed of glycan strands cross-linked by peptides [24]. During cell elongation, the actin homolog MreB controls the spatial pattern of cell-wall synthesis [25, 26] and is essential for maintaining rod-like shape in many bacteria such as *E. coli* [25]. Depletion of MreB [27] or depolymerization of MreB by the small molecule A22 results in cell rounding and eventual lysis [28]. Additionally, sub-inhibitory concentrations of A22 cause cell width to increase and cell length to decrease [29] without affecting cell wall composition [30].

During cell division, cell-wall synthesis localizes to a ring at midcell initialized by the essential and highly conserved tubulin homolog FtsZ [31, 32]. The concentration of FtsZ is negatively correlated with growth rate (and hence cell size) [33–35], and overexpression results in decreased cell length [36]. Moreover, FtsZ overexpression can restore viability to cells depleted of MreB [27], suggesting that FtsZ levels can impact both cell-shape regulation and survival.

Here, we investigated the physiological consequences of Rcs activation in both the absence and presence of cell-envelope damage. We showed that RcsF^IM^ induction reduces growth rate and cell length, increases FtsZ concentration, and reduces the distance between division rings in filamentous cells. Depletion of IgaA resulted in similar phenotypes, indicating that they are general to Rcs activation in the absence of envelope stresses. In fact, cells treated with A22 maintained growth rate and FtsZ concentration even though the Rcs system was activated. Thus, FtsZ levels and cell length are likely downstream of the growth-rate decreases caused by Rcs activation. Altogether, our results show that the nature of an activating stress can affect the phenotypic consequences of a pathway.

## Results

### Induction of a constitutively active RcsF mutant reduces growth rate and cell length

A22 and mecillinam treatment make *E. coli* rounder, increasing cell width and decreasing cell length [29], and activate the Rcs pathway [8]. As we show in an accompanying study, increases in cell width are linked to Rcs activation via altering the periplasmic size [37]. However, in such conditions it is hard to access whether Rcs activation itself affects cellular dimensions and growth rate. To decouple the cue from the response, we reasoned we should control Rcs activation ectopically and used the RcsF mutant RcsF^IM^, which was previously shown to result in constitutive Rcs activation due to its mislocalization to the inner membrane where it alleviates repression of the Rcs signaling by IgaA [8].

To quantify growth behavior and cellular dimensions during exponential growth, we expressed RcsF^IM^ in DH300 Δ*rcsF* cells from a low-copy plasmid (Table S1). We kept cells in steady state and depleted the Rcs proteins produced during stationary phase by continuously diluting RcsF^IM^ cells (Methods). At the same time, we added various concentrations of IPTG and monitored population growth rate, cell size, and Rcs activation levels by beta-galactosidase activity of the promoter of *rprA*, a small RNA expressed when the Rcs pathway is activated [38]. Growth rate decreased and *rprA* expression increased as a function of IPTG concentration, and both saturated at 5-10 µM IPTG (Fig. 1A). Concomitantly, cells decreased in cell length (Fig. 1B). Final yield in stationary phase also decreased with increasing concentrations of IPTG (Fig. S1). Thus, ectopic Rcs activation in the absence of envelope stress slows down growth, decreases cell size, and reduces total biomass production.

**Figure 1:**
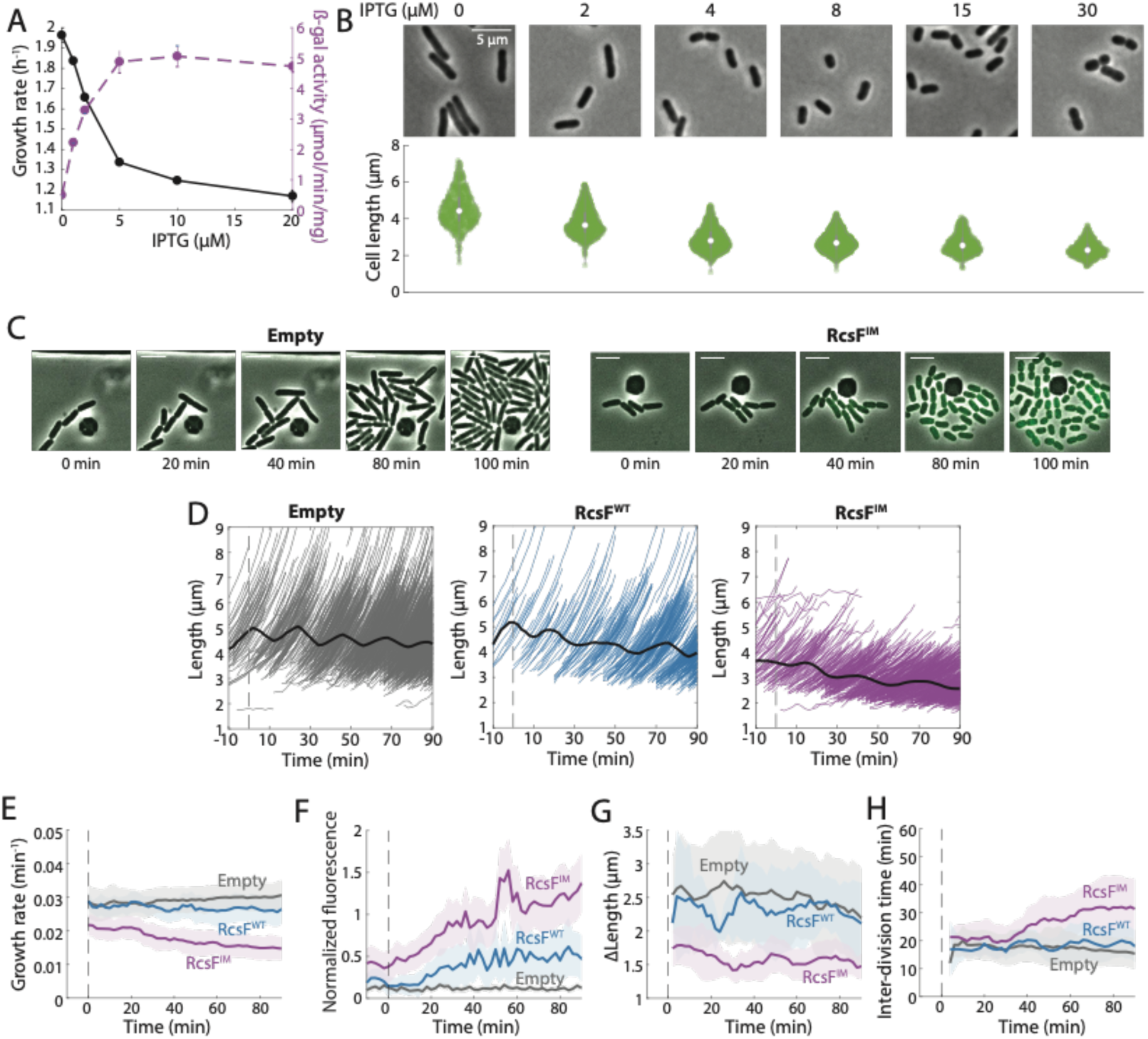
RscF activation in the absence of envelope stress alters cell-length control. A) In batch culture, Δ*rcsF* cells expressing a constitutively active RcsF inner- membrane (RcsF^IM^) variant show a decrease in steady-state growth rate dependent on the level of RcsF^IM^ expression. Error bars represent the standard deviation of three replicate experiments. B) The length of RcsF^IM^-induced cells in batch culture deceased with increasing IPTG concentration. Top: representative cells. Bottom: violin plots of the distribution of cell lengths at each concentration, with *n*>500 cells in each condition. C) After induction of RcsF^IM^ with 15 µM IPTG at *t* = 0, RcsF^IM^ cells decreased in length relative to Δ*rcsF* cells with an empty vector and became physically separated. D) RcsF^IM^ induction at *t*=0 resulted in a gradual reduction in mean cell length (solid black curve, right), while Δ*rcsF* cells (empty vector, left) maintained mean length and RcsF^WT^ resulted in a slight length reduction (middle). Gray, blue, and purple lines are trajectories of individual cells. E) Instantaneous growth rates averaged across the populations in (C) show that RcsF^IM^ induction gradually reduced growth rate, while Δ*rcsF* cells maintained growth rate and RcsF^WT^ resulted in an intermediate growth-rate reduction. F) Mean msfGFP fluorescence averaged across the populations in (C) show that Rcs was activated to a high and intermediate level in RcsF^IM^ and RcsF^WT^ cells, respectively, but not in Δ*rcsF* cells. G) Cell length added over the course of each cell cycle Δ*L* averaged across the populations in (C) stabilized to a lower Δ*L* after induction of RcsF^IM^ compared to Δ*rcsF* or RcsF^WT^-induced cells. H) Division interval averaged across the populations in (C) shows that RcsF^IM^ induction increased the time required for division, but not immediately after induction.

To determine the dynamics of RcsF activation on single cells, and to avoid confounding effects of cell shape changes in optical density measurements, we performed time-lapse imaging in a microfluidic flow cell (Methods) that enabled precise control of induction timing via switching to medium supplemented with IPTG. In this case, we utilized Δ*rcsF* strains with a common plasmid expressing wild-type RcsF (RcsF^WT^) or RcsF^IM^ from an IPTG-inducible promoter (or empty vector), and a second plasmid carrying msfGFP under the *rprA* promoter [38]. Cells were grown for 10 min in LB before 15 µM IPTG was supplemented to reach full Rcs activation by RcsF^IM^ (Fig. 1C). Upon RcsF^IM^ induction, cell length (Fig. 1D) and single-cell instantaneous growth rate (measured as 1/*V dV*/*dt*, where *V* is cell volume, Fig. 1E) decreased steadily, while the empty-vector control strain remained largely unaffected (Fig. 1D,E). The Rcs pathway was fully activated in RcsF^IM^-induced cells (Fig. 1F). RcsF^WT^-induced cells exhibited a small reduction in cell length (Fig. 1D) and growth rate (Fig. 1E), consistent with the observation that RcsF^WT^-induced cells exhibited msfGFP levels intermediate between empty vector and RcsF^IM^ cells (Fig. 1F).

We tracked cell lineages in our time-lapse data to determine whether the volume added during the cell cycle was affected by Rcs activation. Upon IPTG addition, the length added over the course of a cell cycle Δ*L* (a proxy for added volume) in RcsF^IM^ cells equilibrated at 30-40% lower than that of empty-vector or RcsF^WT^ cells (Fig. 1G), consistent with their constant decrease in lower mean cell length (Fig. 1D) and lower growth rate (Fig. 1E). Interestingly, cell-cycle duration increased by 30-40% in 90 min of induction (Fig. 1H). Taken together, these data indicate that Rcs activation alters growth rate, cell size, and cell-cycle timing, motivating us to further investigate the link between the division machinery and Rcs activation.

### Cell shape and growth rate changes are not due to colanic acid production

Production of the exopolysaccharide colanic acid is regulated by the Rcs pathway [39]. In our time-lapse imaging experiments, RcsF^IM^-induced cells became highly separated over time (Fig. 1C,2A), which we hypothesized was due to the production of colanic acid. To determine whether colanic acid production was responsible for the reductions in cell length (Fig. 1D) and growth rate (Fig. 1E) upon Rcs activation, we deleted *wcaJ*, the most upstream gene in the colanic acid synthesis pathway that encodes the initial lipid carrier. Knocking out colanic acid biosynthesis through disruption early in the pathway avoids the buildup of intermediates, which have been shown to change cell shape due to titrating precursor flux away from cell-wall synthesis [40]. We performed time-lapse imaging in a microfluidic flow cell of RcsF^IM^-induced Δ*wcaJ* cells. Δ*wcaJ* cells remained closely packed (Fig. 2A), confirming that the cell separation is indeed due to colanic acid. Moreover, Δ*wcaJ* cells showed a similar decrease in mean cell length (Fig. 2B) and growth rate (Fig. 2C) as colanic acid-producing cells (Fig. 1D,E). Therefore, colanic acid production is not the cause of the cell shape and growth rate changes upon Rcs induction.

**Figure 2:**
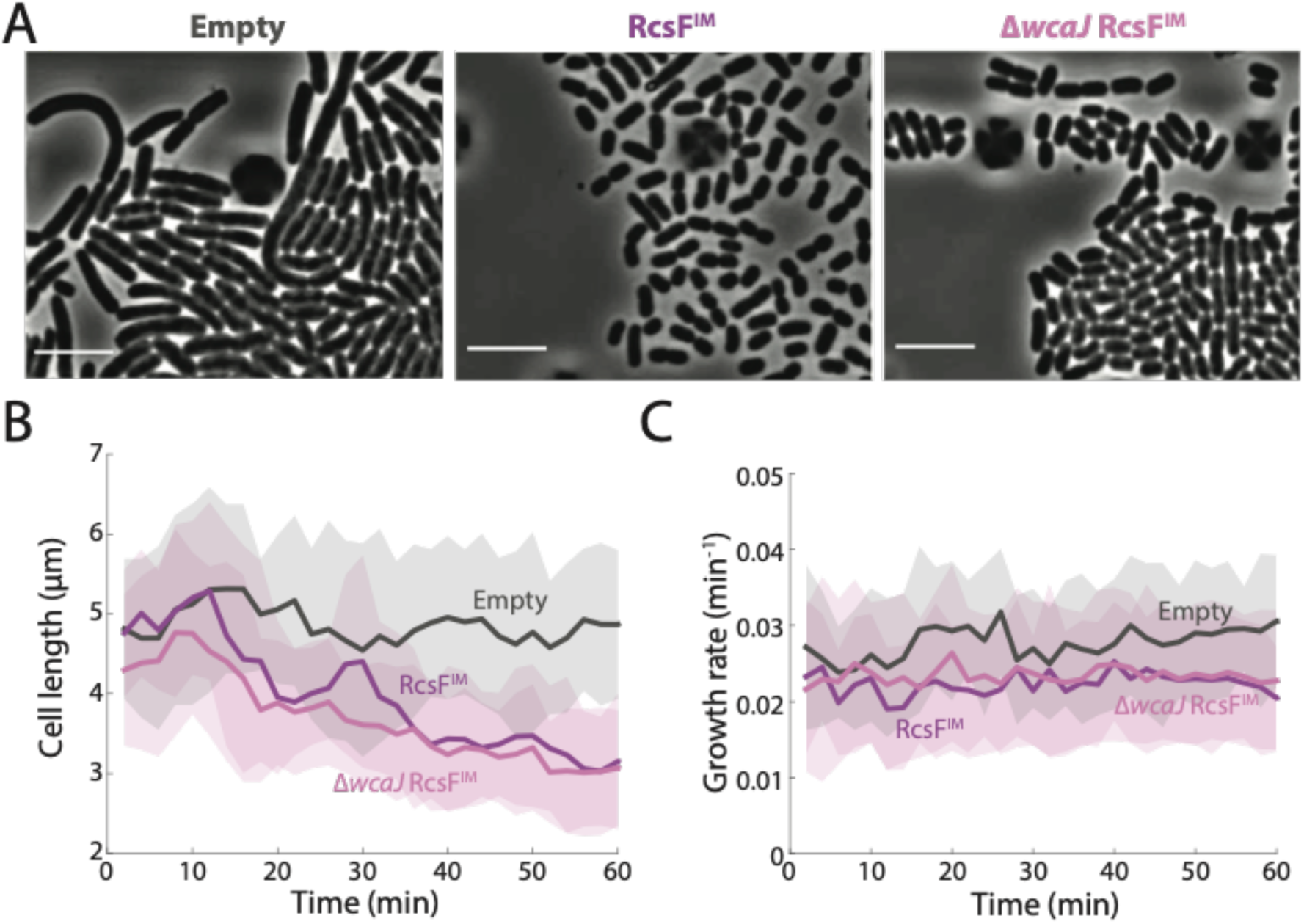
Colanic acid production is not responsible for changes in cell length and growth rate upon RcsF activation. A) Time-lapse of a mutant lacking *wcaJ*, which encodes the most upstream component of colanic-acid synthesis, after IPTG induction of RcsF^IM^ at *t*=0. Unlike wild-type cells (middle), Δ*wcaJ* cells (right) are tightly packed similar to Δ*rcsF* cells with an empty vector (left), which do not activate the Rcs system. B,C) Cell length (B) and growth rate (C) trajectories for Δ*wcaJ* cells are similar to those of RcsF^IM^-induced wild-type cells.

### Constitutive Rcs activation increases FtsZ intracellular concentration and FtsZ localization to division sites

A previous study showed that RcsB, the primary activator of Rcs-regulated genes, can bind upstream of one of the many *ftsZ* promoters, specifically the one that lies also upstream of the preceding *ftsA* [14], and activate *ftsAZ*, suggesting that the Rcs system may regulate expression of the division machinery. To assess this scenario more precisely, we quantified expression of the division machinery upon Rcs induction, and related expression to changes in growth rate and cell length. To quantify total cellular FtsZ concentration as well as the concentration specifically within the FtsZ-ring, we introduced an FtsZ-msfGFP sandwich fusion into the chromosome at the native *ftsZ* locus in *E. coli* DH300 Δ*rcsF* cells with each RcsF plasmid variant. Cells with the FtsZ-msfGFP fusion are viable [41] and have similar growth rate as that of cells with native FtsZ [42]. We imaged cells on agarose pads with M9+0.04% glucose (to alleviate the high autofluorescence of LB) after supplementing them with 0 or 15 µM IPTG. msfGFP intensity (mean total fluorescence normalized to cell volume) was higher in RcsF^IM^-induced compared to uninduced cells (Fig. 3A), indicating that Rcs activation increased FtsZ expression, and RcsF^IM^-induced cells exhibited a ∼50% increase in FtsZ-ring intensity relative to uninduced cells (Fig. 3B). During time-lapse imaging in a microfluidic flow cell, FtsZ-ring intensity increased over time in RcsF^IM^-induced cells, concurrent with a decrease in mean cell length (Fig. 3C). Across a population of single cells, cell length and FtsZ-ring intensity followed a tightly constrained inverse relationship with both 0 and 15 µm IPTG (Fig. 3D). Thus, we inferred that decreases in cell length were coupled to increases in FtsZ ring intensity.

**Figure 3:**
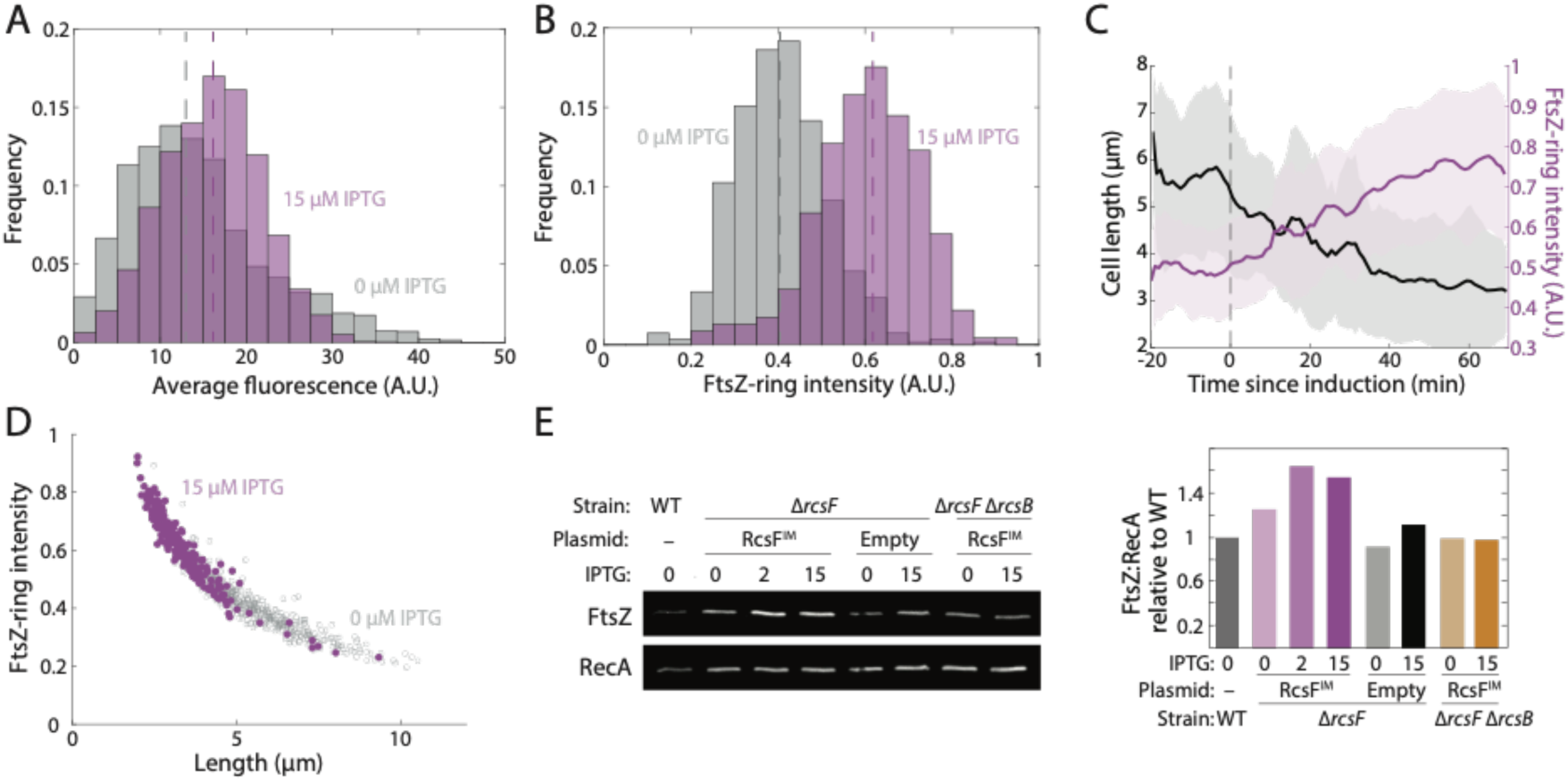
RcsF^IM^ induction results in increased FtsZ concentration within the entire cell and at midcell. A,B) RcsF^IM^ induction with 15 µM IPTG for 60 min increases size-normalized FtsZ-msfGFP fluorescence (A) and FtsZ-ring intensity (B) relative to uninduced cells. Dashed lines represent medians. C) FtsZ-ring intensity (purple) gradually increased as cell length (black) decreased due to RcsF^IM^ induction at *t*=0. D) For both uninduced and induced RcsF^IM^ cells, cell length followed a similar relationship with FtsZ-ring intensity. E) Western blotting demonstrated that the ratio of FtsZ to RecA levels increased dependent on IPTG induction levels in RcsF^IM^ cells but not in cells with an empty vector or cells lacking *rcsB*.

To verify the increase in FtsZ concentration, we directly measured FtsZ protein levels with antibodies. We analyzed wild-type, Δ*rcsF* strains with the empty and RcsF^IM^ plasmids, and a Δ*rcsF* Δ*rcsB* strain with the RcsF^IM^ plasmid (the latter to avoid cell size changes caused by ectopic Rcs induction). Strains were diluted and passaged multiple times to ensure exponential growth, and then treated with 0, 2, or 15 µM IPTG for >90 min. From Western blotting (Fig. 3E), we calculated the ratio of FtsZ to RecA levels (as a control) and normalized to that of wild-type. Basal expression of RcsF^IM^ slightly increased the FtsZ/RecA ratio, and addition of 2 or 15 µM IPTG led to a further increase of ∼50%. This increase required the activation of Rcs-controlled genes by RcsB (Fig. 3E). Thus, Rcs activation leads to an increase in FtsZ levels.

### Rcs induction decreases the separation between FtsZ rings in filamentous cells

Normally dividing cells harbor a single FtsZ ring positioned at midcell, regardless of cell length [33]. To determine whether FtsZ localization dynamics and local variations in FtsZ concentration were affected by RcsF^IM^ induction and connected with reductions in cell length, we treated cells with cephalexin, a beta-lactam that inhibits the division-specific transpeptidase FtsI [43]. These filamentous cells had multiple FtsZ rings that were easily identified based on the peaks in peripheral fluorescence (Methods) [42]. After growth on LB agarose pads with 15 µM IPTG, kymographs indicated that RcsF^IM^-induced cells had more closely spaced FtsZ-rings than empty-vector cells (Fig. 4A). Indeed, RcsF^IM^-induced cells exhibited an increased number of rings per unit length (Fig. 4B) and decreased distance between FtsZ rings (Fig. 4C) compared with empty- vector or RcsF^WT^-induced cells. Thus, RcsF^IM^ induction increases the number of FtsZ rings per unit length, thereby defining shorter cellular units even in the absence of cytokinesis.

**Figure 4:**
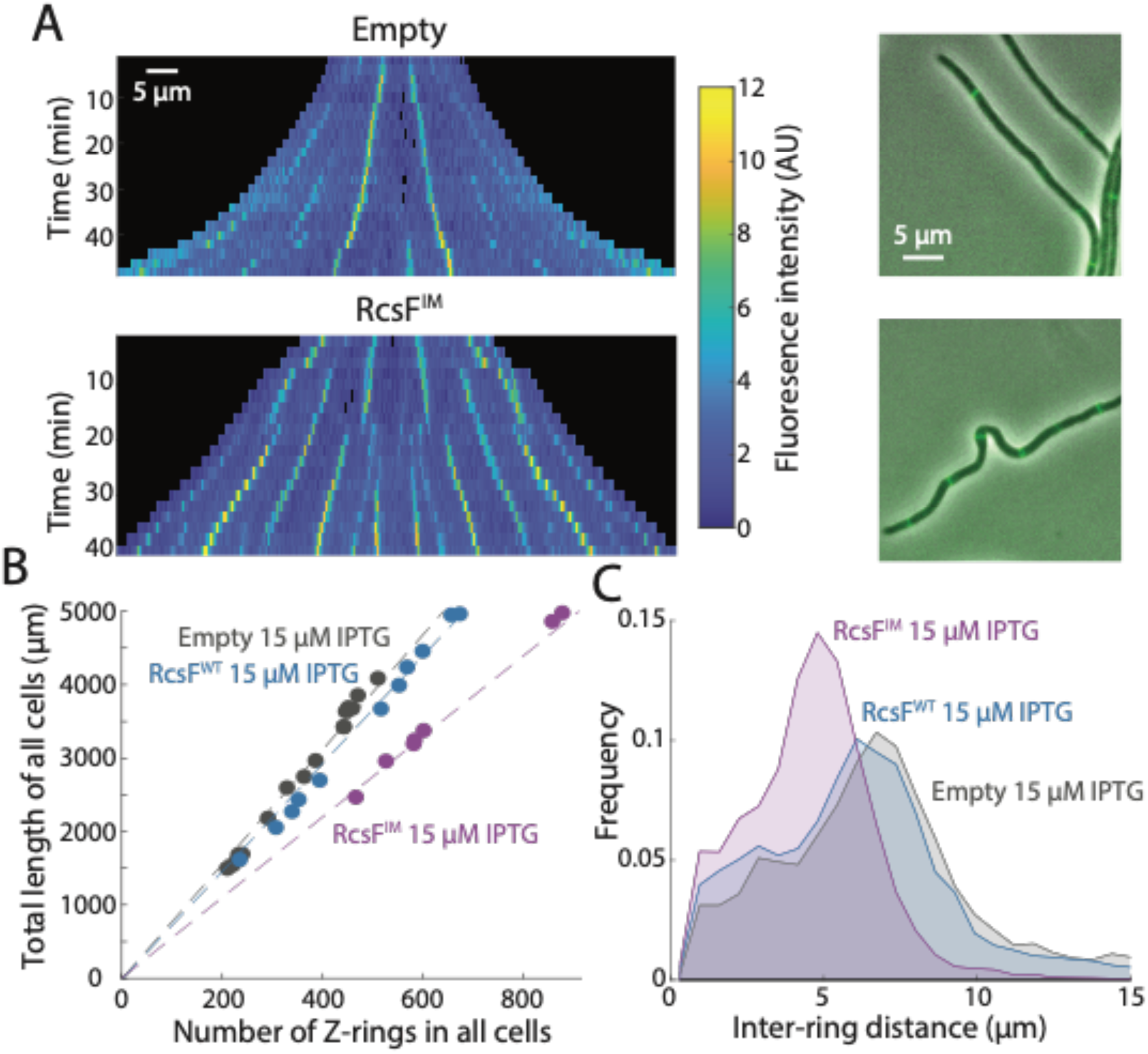
RcsF^IM^ induction decreases the distance between FtsZ rings in filamentous cells. A) After cephalexin treatment to inhibit division, kymographs (left) and overlays of FtsZ-msfGFP fluorescence and phase-contrast images at *t*=40 min (right) during induction with 15 µM IPTG show that the distance between FtsZ rings is shorter in RcsF^IM^-induced cells (bottom) compared to empty-vector Δ*rcsF* cells (top). B) The slope of the total length of a population of cells versus the number of Z-rings within those cells is lower in RcsF^IM^-induced cells. Each circle represents all cells within a field of view. C) The distance between FtsZ-rings is significantly shorter in RcsF^IM^ cells compared to RcsF^WT^ cells and empty-vector Δ*rcsF* cells.

### IgaA depletion mimics RcsF^IM^ induction

To determine whether the changes in growth rate (Fig. 1E), cell length (Fig. 1D), and FtsZ concentration (Fig. 3) were general features of Rcs activation as opposed to an artefact of the RcsF^IM^ mutant, we decided to use a different way to induce the Rcs system, this time using the essential protein IgaA, which is an inner membrane inhibitor of the Rcs system [10]. We utilized a strain expressing the *igaA^L523A^* allele from an arabinose-inducible promoter, with the native *igaA* deleted. This strain, akin to the previously used *igaA^L643P^* allele [8, 44], exhibits lower levels of Rcs repression and hence the effects of depletion are observed more rapidly than depletion of wild-type IgaA. To determine whether IgaA^L523A^ depletion and the consequent activation of the Rcs system affects FtsZ concentration, we transduced the FtsZ-msfGFP sandwich fusion onto the chromosome of the *igaA^L523A^*-inducible strain; all measurements were performed in strains with the FtsZ-msfGFP sandwich fusion. We monitored growth of the strain via absorbance in LB supplemented with either 0.2% arabinose, to maintain normal growth, or 0.2% glucose, to deplete IgaA^L523A^. With arabinose, the IgaA^L523A^ strain grew similarly to wild-type *E. coli*, while depletion with glucose substantially reduced growth rate (1.76±0.17 h^-1^ in arabinose versus 1.15±0.20 h^-1^ in glucose; *p* < 10^-7^, *t*-test) (Fig. 5A, S2), similar to the decrease in growth rate upon RcsF^IM^ induction (Fig. 1A).

**Figure 5:**
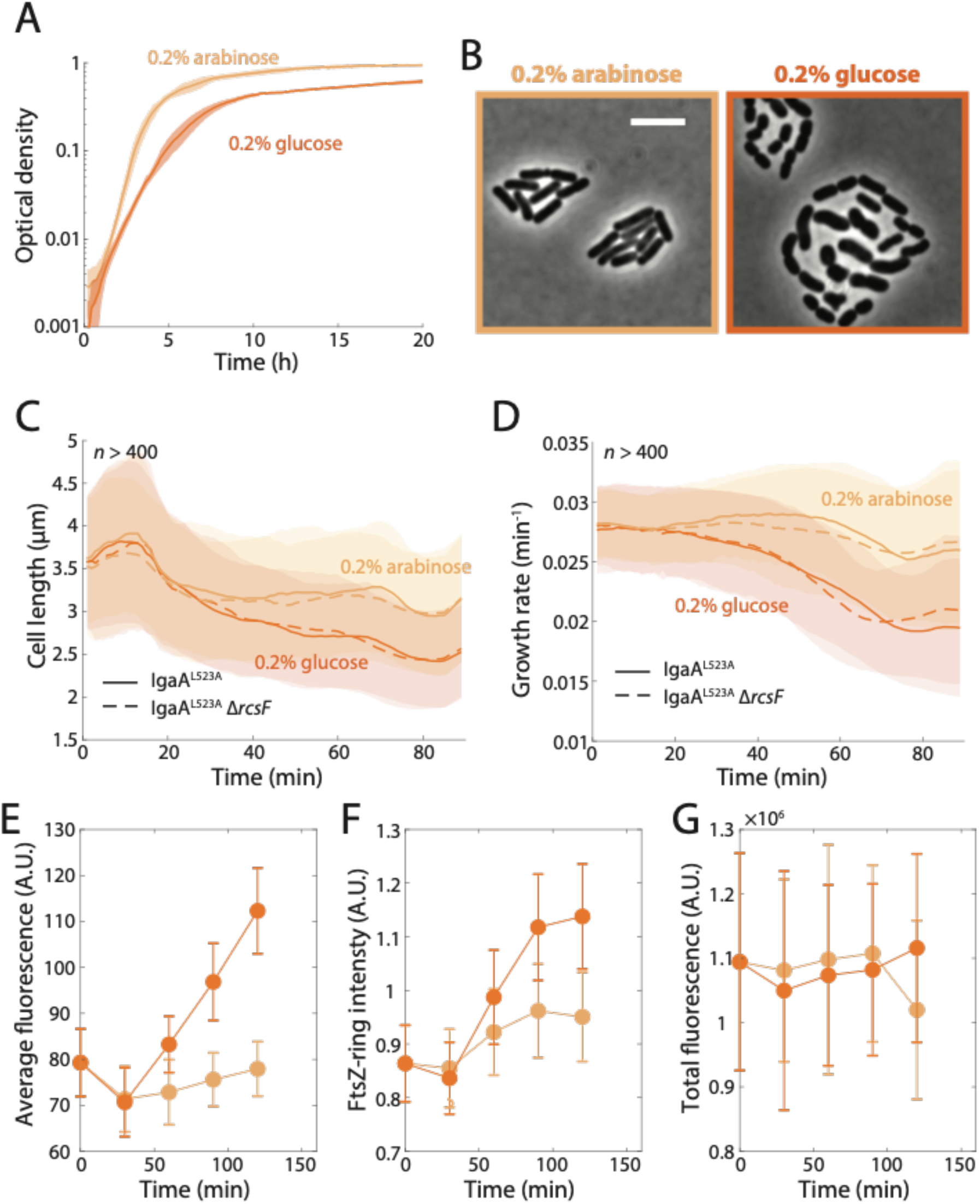
IgaA^L523A^ depletion results in similar phenotypes to RcsF^IM^ induction. A) Using a strain with a plasmid expressing the *igaA^L523A^* allele from an arabinose-inducible promoter with the native *igaA* deleted (KC1183, Table S1), depleting IgaA^L523A^ with glucose results in a reduction in growth. Shaded areas represent 1 standard deviation (S.D.) across *n*=12 replicates. B) IgaA^L523A^ depletion (glucose) resulted in shorter cells that became physically separated, indicating that the Rcs system is activated. Images are 60 min after imaging. C,D) IgaA^L523A^ depletion (glucose) caused cell length (C) and growth rate (D) to decrease, independent of RcsF, which lies upstream of IgaA in the Rcs signaling cascade. Comparing data with (KC1183) and without *rcsF* (KC1185, Table S1) between 60-80 min, length (glucose, *p*=0.34; arabinose, *p*=0.12) and growth rate (glucose, *p*=0.61; arabinose, *p*=0.31) were not significantly different, while *p*<10^-10^ for all comparisons between glucose and arabinose. *p-*values are from two-sided Student’s *t*-tests with *n*>500 cells in each condition. Data points are mean±S.D. E-G) In strains expressing an FtsZ-msfGFP sandwich fusion, average fluorescence (E) and FtsZ-ring intensity (F) increased upon IgaA depletion. Total fluorescence (G) remained similar in arabinose and glucose. Data points are mean±S.D., with *n*>50 cells each condition.

We next examined the IgaA^L523A^ strain during depletion using single-cell microscopy. When spotted onto agarose pads with 0.2% glucose, cells physically separated during growth and division, signifying colanic acid production due to Rcs activation (Fig. 5B). Mean cell length (Fig. 5C) and instantaneous growth rate (Fig. 5D) decreased in a manner similar to RcsF^IM^ induction (Fig. 1C,D). Size-normalized FtsZ-msfGFP fluorescence intensity throughout the cell (Fig. 5E) and FtsZ-ring intensity (Fig. 5F) increased coincident with the decrease in length (Fig. 5C), such that total fluorescence remained approximately constant (Fig. 5G). In sum, the phenotypes that emerged during RcsF^IM^ induction were closely mimicked by IgaA^L523A^ depletion, suggesting that they are general properties of Rcs activation in the absence of envelope stress.

### A22 activates RcsF without affecting growth rate or FtsZ concentration

To investigate the role of the Rcs system in the response to A22 at the cellular level, we used time-lapse microscopy to analyze the response of wild-type cells to a range of A22 concentrations (Fig. 6A). As expected, mean cell width increased upon exposure to A22 in a dose-dependent manner (Fig. 6B) [30].

**Figure 6:**
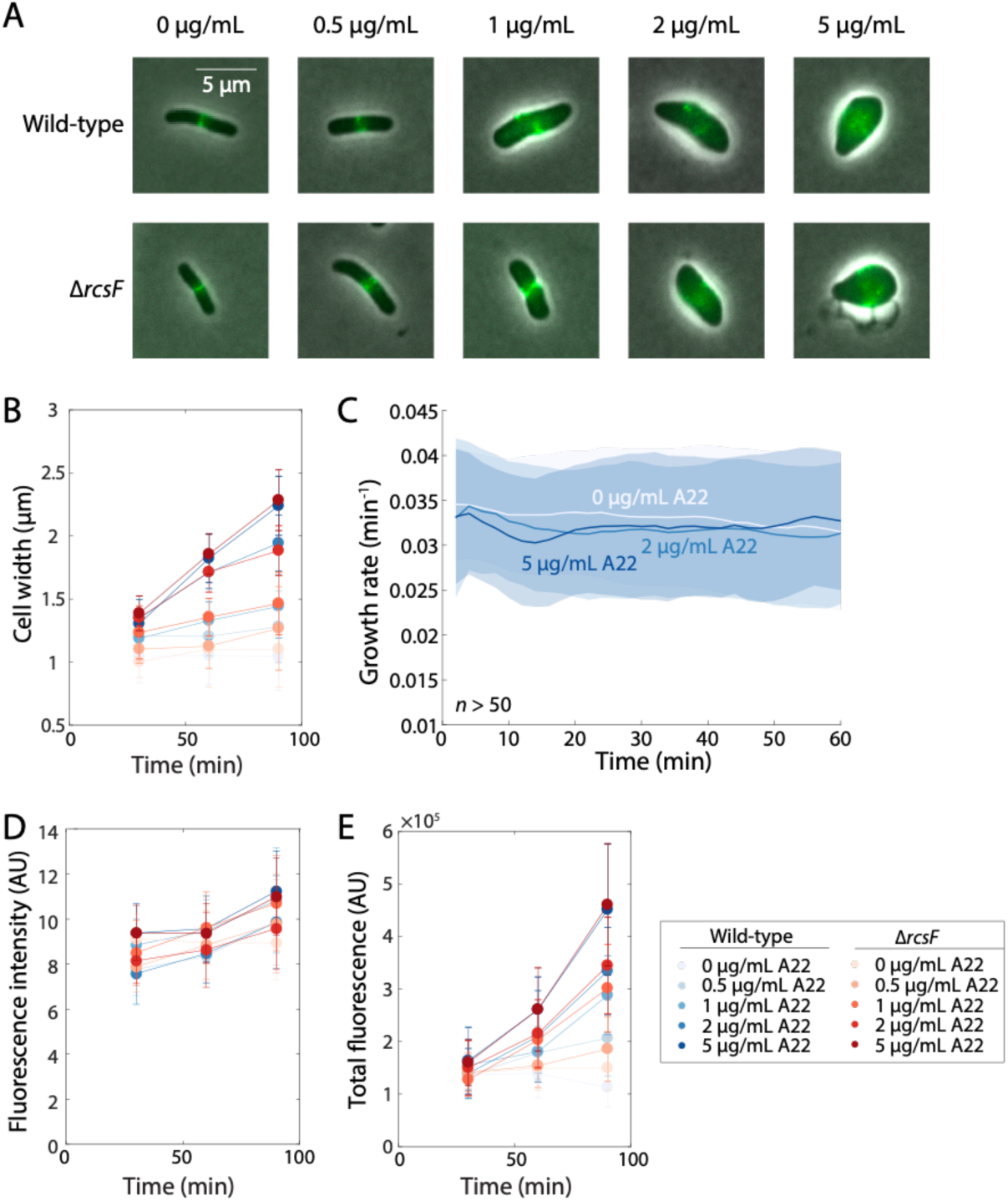
A22 treatment does not result in changes to growth rate or FtsZ concentration. A) A22 treatment caused the width of both wild-type and Δ*rcsF* cells to increase. FtsZ-msfGFP fluorescence is overlaid on phase-constrast images. Images were taken after 90 min with A22 treatment. B) Cell width increased in an A22 dose-dependent manner, with similar increases in wild-type and Δ*rcsF* cells. A22 was added at *t*=0. *n*>50 cells per condition. C) Log-phase wild-type cells in LB were placed onto LB-agarose pads containing different concentrations of A22, and their growth was monitored by time-lapse imaging. Throughout the experiment, growth rate was unaffected by A22 treatment. D) A22 was added to log-phase wild-type or Δ*rcsF* cells in liquid LB cultures at *t*=0. Every 30 min, small aliquots of cells were imaged on agarose pads to monitor changes to cell morphology and FtsZ intracellular intensity (total fluorescence normalized by cell area). FtsZ-msfGFP fluorescence intensity in wild-type and Δ*rcsF* cells increased only slightly or not at all and was similar across all A22 concentrations. E) Total FtsZ-msfGFP fluorescence in wild-type and Δ*rcsF* cells increased over time, reflecting the increase in cell width and volume (B) and maintenance of FtsZ-msfGFP concentration (D). Data points in (B,D,E) are mean±1 standard deviation. *n* > 50 cells were analyzed for each condition.

However, despite rapid activation of the Rcs system by A22 [37], growth rate was essentially maintained for the first hour (Fig. 6C), consistent with previous studies showing that A22 does not affect growth rate [45].

This growth rate maintenance is markedly distinct from other Rcs-activating perturbations such as RcsF^IM^ induction (Fig. 1D) or IgaA^L523A^ depletion (Fig. 5D). Moreover, there was little increase in the volume-normalized FtsZ-msfGFP intensity (Fig. 6D), as both total fluorescence and cell width increased together: ∼3-fold for the former (Fig. 6E) and ∼75% for the latter (Fig. 6B) at the highest A22 concentration. These phenotypes are also distinct from the increase in FtsZ concentration (Fig. 3A,E, 5E) observed for Rcs activation in the absence of envelope stresses. Thus, surprisingly A22 somehow reverses the growth inhibition that normally occurs under Rcs activation, and prevents upregulation of FtsZ concentration.

Consistent with the maintenance of FtsZ concentration (Fig. 6D) despite activation of the Rcs system under A22 treatment [37], the trajectories of cell width (Fig. 6B) and growth rate (Fig. 6C) were quantitatively similar in A22- treated wild-type and Δ*rcsF* cells, indicating that the morphological effects of A22 treatment (cell widening) are independent of the Rcs pathway. Consistent with this finding, wild-type and Δ*rcsF* cells exhibited similar shape trajectories during recovery from A22 (Fig. S3). Given that FtsZ concentration varies with nutrient- determined growth rate [33], these data suggest that the dynamics of cellular dimensions and FtsZ are downstream of the growth-rate effects of Rcs activation.

## Discussion

In this study, we showed that activation of the Rcs system in the absence of envelope stress adjusts the added length (τ−*L*) per cycle and decouples elongation and cell-cycling timing, leading to shorter cells (Fig. 1C-F). This phenotype is achieved by up-regulation of FtsZ levels and FtsZ ring formation (Fig. 3A,B,E), which likely are consequences of the slowdown in growth due to Rcs activation [33]. Although the up-regulation in FtsZ protein levels may be due in part to the effect of Rcs on *ftsAZ* transcription [14], how FtsZ increases its mid-cell localization remains to be elucidated. Overall, our data highlight the strong coupling between growth rate and FtsZ levels, the latter of which determines cell length in a highly stereotyped manner (Fig. 3D).

The growth-rate decrease and upregulation of FtsZ in RcsF^IM^-induced cells (Fig. 1D,E) is a direct result of Rcs activation, as IgaA depletion had similar effects (Fig. 5D,E). Thus, these phenotypes are likely to be general consequences of Rcs activation in the absence of envelope stresses. However, A22 treatment did not affect growth rate (Fig. 6C) or FtsZ concentration (Fig. 6D) despite activation of the Rcs pathway [37]. Moreover, A22 treatment resulted in cell-length decreases similar to RcsF^IM^ induction even in the absence of RcsF (Fig. S4), indicating that there are Rcs-independent mechanisms for coupling the cell cycle to cellular dimensions [17, 46], and/or there may be additional signals that buffer or alter the Rcs response during cell elongation stress. Together, these findings indicate that the results of activating a stress response can be highly dependent on context. They also provide a plausible explanation for why *igaA* deletions are non-viable in enterobacteria, unless Rcs is also knocked out [8, 44]. As we have shown, the resulting constant activation of the Rcs pathway leads to cells becoming very small, likely tuning ϕ..*L* below a level that supports growth. This ability to transition to a non-growing state may be used by enterobacteria to enter a persistent state in a host, as IgaA is one of the most down-regulated proteins in non-dividing *Salmonella* cells in host tissues [47]. Interestingly, LPS-related stress that induces Rcs independently of cell size cues [37] causes transient Rcs activation that fixes the damage [48, 49].

The relationship between the Rcs pathway and cell division has long been the subject of speculation, with overexpression of RcsB and RcsF previously shown to suppress the division defect of an *ftsZ84* mutant [50, 51]. Further analysis of the FtsZ promoter sequence identified an RcsB binding site in *ftsAZ* that enables transcription of *ftsZ* upon Rcs induction [14]. Our findings present a more nuanced perspective on Rcs-mediated regulation of the division machinery, in which increases in FtsZ concentration occur only for conditions in which growth rate decreases, such as RcsF^IM^ induction (Fig. 1D) or IgaA depletion (Fig. 5D), but not A22 treatment (Fig. 6C). Taken together, our findings reinforce previous studies linking the elongation and division machineries [42, 52, 53] and highlight the importance of growth rate in determining cell size.

Understanding how the Rcs system enacts physiological changes in bacterial cells has important implications for the general response of bacteria to stresses. One significant takeaway from our study is that cell-shape and transcriptional changes can result from a global change in growth rate rather than directly from stress-response pathway activation. That cells maintain growth rate during A22 treatment in the face of Rcs activation suggests that other responses induced by A22 interfere with some of the downstream Rcs signaling. It will be interesting to probe the extent to which activation of stress response pathways generally lead to changes in growth rate; our findings predict that activation of growth-inhibiting pathways will also cause *ftsZ* upregulation. Moreover, changes in growth rate could lead to cross protection of other stresses [54]; for example, growth in minimal medium alleviates the essentiality of MreB [55]. Ultimately, our strategy of decoupling Rcs activation from the activating stresses should be a powerful strategy for connecting response pathways to their ensuing the physiological phenotypes.

## Methods

### Strains, plasmids, and media

All strains and plasmids are in Table S1. *E. coli* MG1655 and derivatives of it were used in all experiments. We used P1 transduction to introduce *ftsZ-msfGFP* into the chromosome at the native *ftsZ* locus of DH300 Δ*rcsF* cells, and introduced the pNG162-empty, pNG162-WT, and pNG162-IM plasmids harboring the RcsF variants, along with the pTrcHis2A plasmid, which carries *lacI*^q^ from a high copy plasmid to shut down basal expression from the pNG162 plasmid [8].

Cells were normally cultivated in LB Lennox (10 g/L tryptone, 5 g/L yeast extract, 5 g/L NaCl), or EZ-RDM [56] or M9 [57] media for imaging. Antibiotics were added to the culture when needed to maintain the plasmid.

### Growth-rate measurements for batch cultures

For Fig. 1A and S1, overnight cultures were grown at 37 °C in LB (Lennox formulation) with appropriate antibiotics. Cells were diluted to OD578=0.001, and grown for ∼3 h until OD578∼0.1. Cells were then re-diluted to OD578=0.025 in fresh LB with 0 to 15 µM IPTG. To enable growth-rate quantification, OD578 was measured throughout the experiment, and once OD578=0.3 was reached, cells were diluted to OD578=0.025. This passaging cycle was maintained for 6 to 9 h. Growth rate was calculated from OD578 measurements using ≥3 measurements from the range with OD578<0.4 by fitting to an exponential.

For the measurements in Fig. 5A, overnight cultures in LB with 100 µg/mL and 0.2% arabinose were inoculated into 200 µL of fresh media supplemented with 100 µg/mL of ampicillin and 0.2% of arabinose or glucose in a clear 96-well plate. The plate was covered with an optical film, with small holes poked at the side of each well to allow aeration. Incubation and OD measurements were performed with an Epoch 2 plate reader (BioTek) at appropriate temperatures with continuous shaking and OD600 measured at 7.5-min intervals. Growth rate was calculated as the slope of ln(OD) with respect to time after smoothing using a moving average filter of window size five.

### Imaging acquisition on agarose pads

For fixed time-point experiments, cells were diluted 1:5000 from an overnight culture. For experiments in Fig. 3A,B, Fig. 4B,C, and Fig. 5E-G, diluted cells were grown for 3 h to OD∼0.1, then diluted 1:10 into fresh medium with appropriate inducers. For experiments in Fig. 6A,B,D,E, S3, and S4, diluted cells were grown to OD=0.4, then diluted 1:200 in 0, 0.25, 0.5, 1, 2, or 5 µg/ml A22. In these experiments, a small aliquot of cells (∼ 1 µL) was placed onto agarose pads with M9+0.04% glucose every 30 min. Phase contrast and GFP fluorescence images were acquired as quickly as possible to avoid cell shape changes due to the medium change.

For time-lapse experiments, cells were diluted 1:5000 from an overnight culture. For experiments in Fig. 3C, 4A, and 5B-D, after 3 h of the 1:5000 dilution, cells were further diluted 1:10 and spotted onto EZ-RDM pads (Fig. 3C), LB pads with 8 µM IPTG and 10 µg/mL cephalexin (Fig. 4A), or LB pads with 0.2% arabinose/glucose (Fig. 5B-D). For experiment in Fig. 6C, at OD=0.2, cells were diluted 1:10 onto LB pads with 1% agarose and 0, 0.25, 0.5, 1, 2, or 5 µg/ml A22.

Cells were imaged under phase contrast and fluorescence. The agarose pads were maintained at 37 °C using a heated environmental chamber (Haison Tech). Phase contrast and fluorescence images were acquired every 2 min. All phase contrast and fluorescence images were collected on a Nikon Ti-E epifluorescence microscope using µManager v. 1.4 [58].

### Imaging in microfluidic flow cells

For imaging experiments in Fig. 1C-H and Fig. 2, cells were diluted 1:500 from overnight cultures and grown for 3.5 h. Cells were then diluted to approximately OD=0.001 and placed in a CellAsic flow cell chamber pre-warmed to 37 °C. Once introduced into the imaging chamber, cells were grown in LB for 20 min before switching to LB supplemented with IPTG.

### Quantification of FtsZ fluorescence levels and spatial distribution

All phage contrast images were segmented using *Morphometrics* [26]. FtsZ rings were detected based on peaks in intensity along the contour. FtsZ ring intensity was calculated as the difference between the maximum and minimum of the fluorescence peak times the width of the peak, after background subtraction [42]. Fluorescence intensity was calculated as the sum of all pixels in the fluorescence channel within the contour of the cell divided by the area of contour, after background subtraction. For data in Fig. 4B, FtsZ ring distances were calculated based on adjacent fluorescence peaks as extracted from the midline of the cells.

### Western blot quantification

After incubation with IPTG for at least 1.5 h, 1 mL of exponentially growing culture was harvested, and lysed and solubilized by boiling in Laemmli buffer for 5 min at 95 °C. Samples were diluted and normalized based on OD578 prior to SDS-PAGE analysis. Proteins were separated on a 15% SDS-PAGE gel, and transferred on PVDF membranes (Immobilon-P). Membranes were blocked with 5% skim milk in TBS-T (50 mM Tris-HCl [pH 7.6], 0.15 M NaCl, and 0.1% Tween20). TBS-T was used in all subsequent steps of the immunoblotting procedure. Anti-FtsZ (1:1000, Acris) and anti-RecA (loading control, 1:1000, Abcam) rabbit antisera were used as primary antibodies. Membranes were incubated with secondary antibodies conjugated with horseradish peroxidase (HRP) diluted in 5% skim milk in TBS-T (1:10,000, GE healthcare). Labelled proteins were detected via enhanced chemiluminescence (Pierce ECL Western Blotting Substrate, Thermo Scientific) and exposed on X-ray films (Kodak Biomax Mr1).

The built-in gel analysis tool in FIJI [59] was used to quantify FtsZ and RecA levels from a horizontal rectangle including the relevant bands.

## Acknowledgements

The authors thank the Huang and Typas labs for useful discussions. This work was supported by a National Science Foundation Graduate Research Fellowship (to A.M.), an ARCS Fellowship (to A.M.), a James McDonnell Postdoctoral Fellowship (to H.S.), EMBL core funding and a DFG grant (TY 116/2-1) for SPP1617 (to A.T.), NIH Director’s New Innovator Award DP2OD006466 (to K.C.H.), NSF CAREER Award MCB-1149328 and grant EF-2125383 (to K.C.H.), and the Allen Discovery Center at Stanford on Systems Modeling of Infection (to K.C.H.). K.C.H. is a Chan Zuckerberg Biohub Investigator. This work was also supported in part by the National Science Foundation under Grant PHYS- 1066293 and the hospitality of the Aspen Center for Physics.

## Supplemental Figures

**Figure S1:**
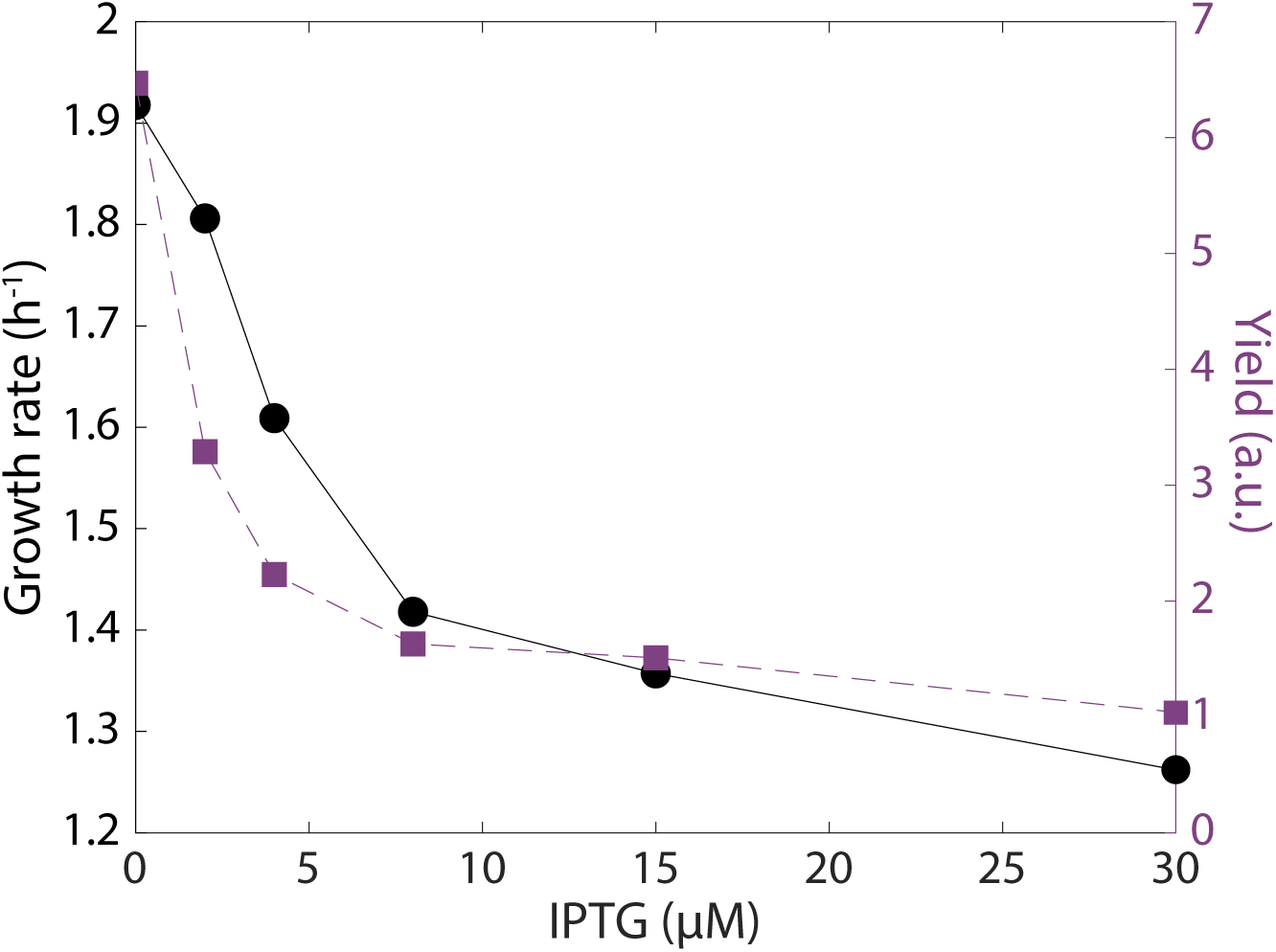
RcsFIM induction led to a decrease in steady-state growth rate and stationary-phase yield in batch culture.

**Figure S2:**
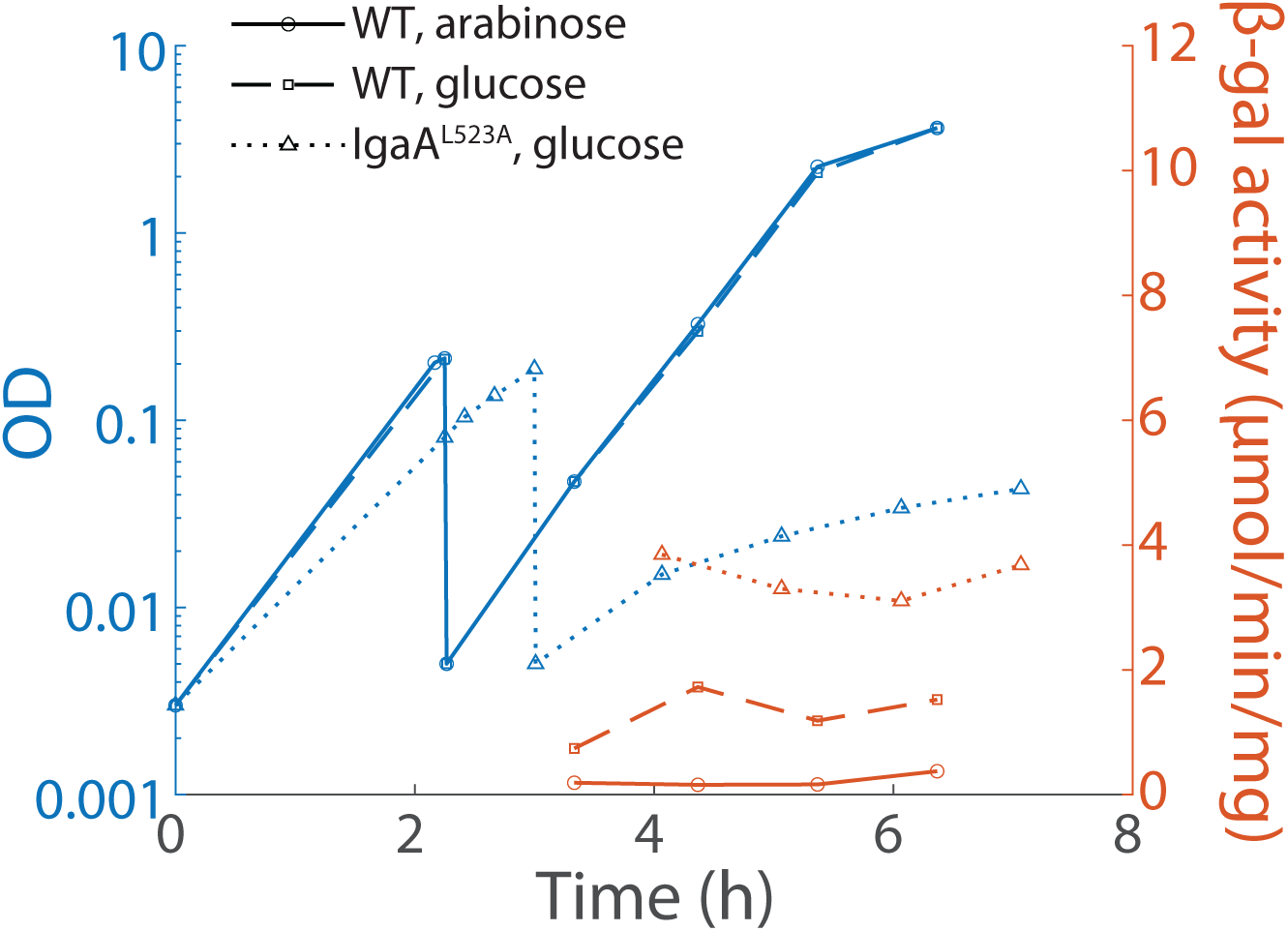
Depletion of IgaA^L523A^ decreases growth rate and activates the Rcs system. Activation was measured through induction of chromosomal *rprA::lacZ* in a β-galactosidase assay (right). IgaA^L523A^ cells were switched to glucose at *t*=0.

**Figure S3:**
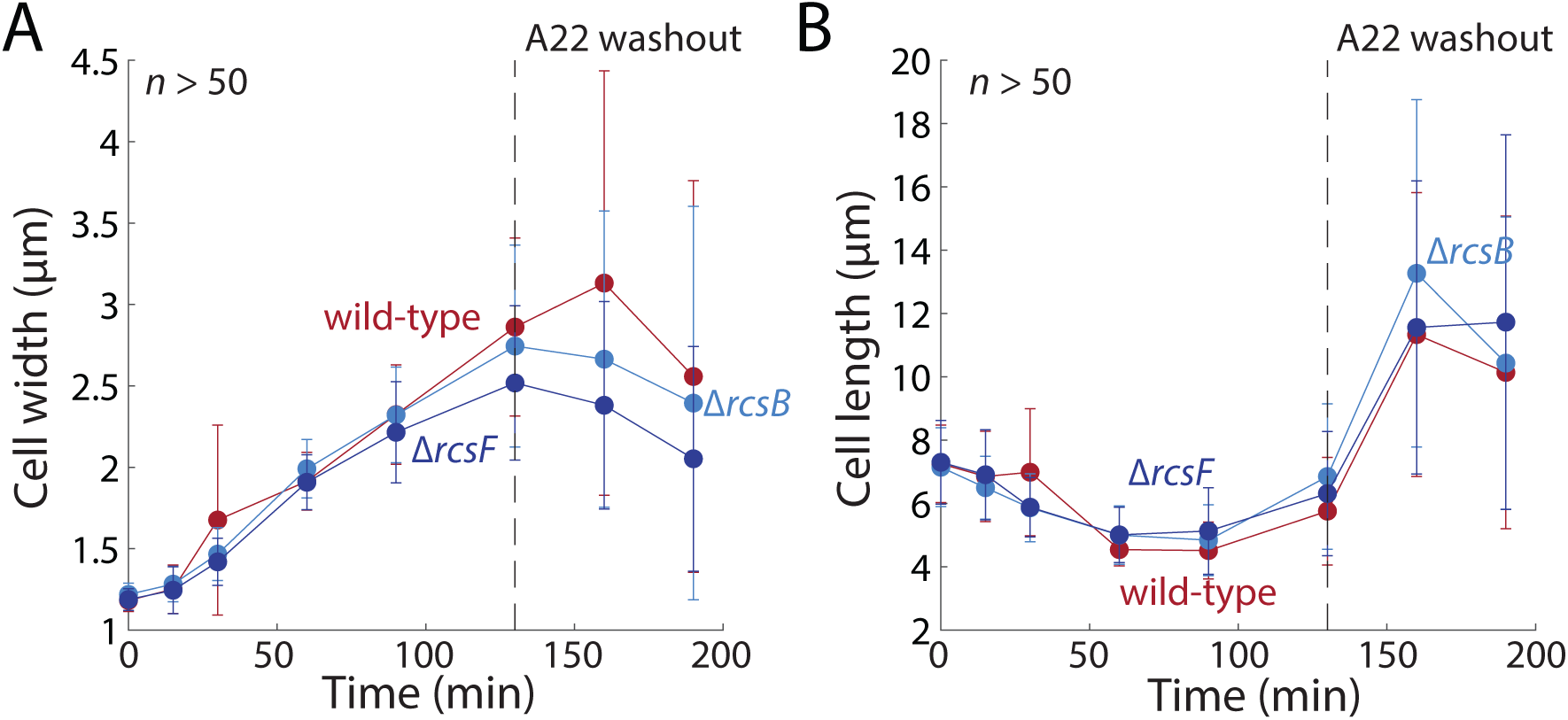
Cell shape recovery after A22 treatment is Rcs-independent. Cells growing in LB were treated with 5 µg/mL A22 at *t*=0 and aliquots of the cultures were sampled periodically to measure the dynamics of width (A) and length (B). After 130 min, cells were resuspended in LB without A22. Similar dynamics were observed in wild-type, Δ*rcsF*, and Δ*rcsB* cells. Data points are mean±1 standard deviation with *n*>50 cells.

**Figure S4:**
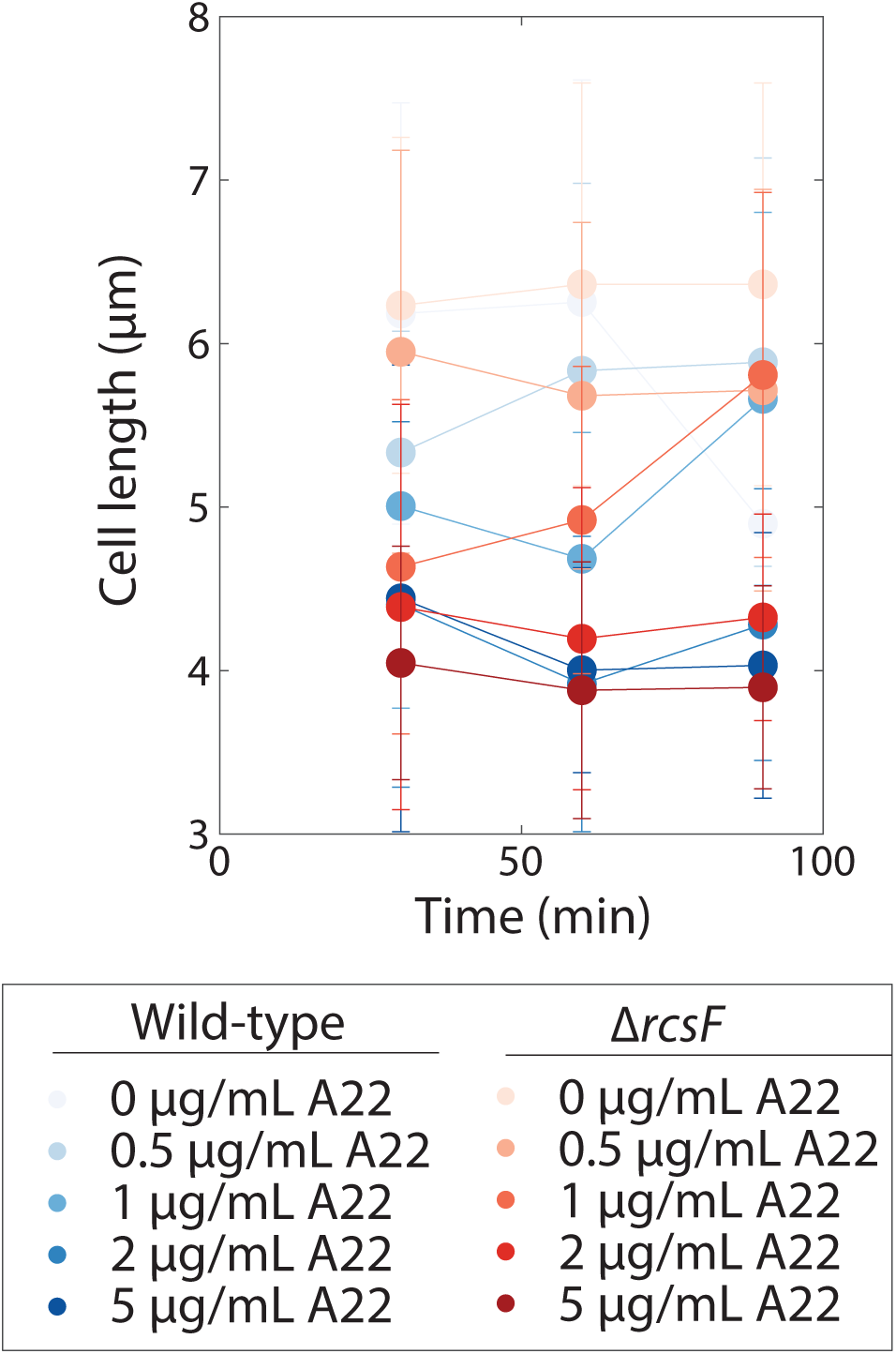
Cell length decreases in an A22 dose-dependent but RcsF- independent manner. Cells are the same as in Fig. 6. Data are mean±1 standard deviation with *n*>50 cells.

## Supplemental Tables

**Table S1:** Strains used in this study.

| Strain | Relevant genotype or features | Source |
| --- | --- | --- |
| DH300 | *rprA::lacZ* MG1655 (*argF-lac*)U169 (used as wild type) | [38] |
| NT15048 | DH300 $\Delta rcsF::kan$ | This study |
| AM110 | DH300 $\Delta rcsF::cat \Delta wcaJ::kan$ | This study |
| NT15178 | DH300 $\Delta rcsF::kan \Delta rcsB::cm$ | This study |
| AM115 | DH300 $\Delta rcsF ftsZ::ftsZ-msfGFP$ | [41], this study |
| KC1183 | DH300 pBAD30-IgaA<sup>L523A</sup> $\Delta igaA::cat$ | This study |
| KC1185 | DH300 $\Delta rcsF::kan$ pBAD30-IgaA<sup>L523A</sup> $\Delta igaA::cat$ | This study |
| AM181 | DH300 pBAD30-IgaA<sup>L523A</sup> $\Delta igaA::cat ftsZ::ftsZ-msfGFP$ | This study |

**Table S2:** Plasmids used in this.

| Plasmid | Features | Source or notes |
| --- | --- | --- |
| pAC581 |  | [60] |
| pBAD30-IgaA<sup>L523A</sup> | IgaA<sup>L523A</sup> under arabinose-induced promoter, ampicillin | This study |
| pMZ13 | msfGFP under *rprA* promoter, pAC581-based, chloramphenicol | This study |
| pNG162-Empty | Control plasmid, spectinomycin | [8] |
| pNG162-RcsF-WT | RcsF<sup>WT</sup> under IPTG-induced promoter, spectinomycin | [8] |
| pNG162-RcsF-IM | RcsF<sup>IM</sup> under IPTG-induced promoter, spectinomycin | This study |
| pTrcHis2A | ampicillin | Invitrogen |

## References

1. Jonas K. To divide or not to divide: control of the bacterial cell cycle by environmental cues. Current opinion in microbiology. 2014;18:54–60.

2. Schaechter M, Maaloe O, Kjeldgaard NO. Dependency on medium and temperature of cell size and chemical composition during balanced grown of Salmonella typhimurium. Journal of general microbiology. 1958;19(3):592–606. PubMed PMID: 13611202.

3. Hengge-Aronis R. Signal transduction and regulatory mechanisms involved in control of the σS (RpoS) subunit of RNA polymerase. Microbiol Mol Biol Rev. 2002;66(3):373–95.

4. Matin A. The molecular basis of carbon-starvation-induced general resistance in Escherichia coli. Molecular microbiology. 1991;5(1):3–10.

5. Majdalani N, Gottesman S. The Rcs phosphorelay: a complex signal transduction system*. Annual Review of Microbiology. 2005;59(1):379–405. doi: 10.1146/annurev.micro.59.050405.101230. PubMed PMID: 16252004676231035489related:Yfq8biW4iuEJ.

6. Wall E, Majdalani N, Gottesman S. The Complex Rcs Regulatory Cascade. Annu Rev Microbiol. 2018;72:111–39. Epub 2018/06/14. doi: 10.1146/annurev-micro-090817-062640. PubMed PMID: 29897834.

7. Nichols RJ, Sen S, Choo YJ, Beltrao P, Zietek M, Chaba R, et al. Phenotypic landscape of a bacterial cell. Cell. 2011;144(1):143–56. Epub 2010/12/28. doi: 10.1016/j.cell.2010.11.052. PubMed PMID: 21185072; PubMed Central PMCID: PMCPMC3060659.

8. Cho SH, Szewczyk J, Pesavento C, Zietek M, Banzhaf M, Roszczenko P, et al. Detecting envelope stress by monitoring beta-barrel assembly. Cell. 2014;159(7):1652–64. doi: 10.1016/j.cell.2014.11.045. PubMed PMID: 25525882.

9. Konovalova A, Perlman DH, Cowles CE, Silhavy TJ. Transmembrane domain of surface-exposed outer membrane lipoprotein RcsF is threaded through the lumen of beta-barrel proteins. Proc Natl Acad Sci U S A. 2014;111(41):E4350–8. Epub 20140929. doi: 10.1073/pnas.1417138111. PubMed PMID: 25267629; PubMed Central PMCID: PMCPMC4205638.

10. Hussein NA, Cho SH, Laloux G, Siam R, Collet JF. Distinct domains of Escherichia coli IgaA connect envelope stress sensing and down-regulation of the Rcs phosphorelay across subcellular compartments. PLoS Genet. 2018;14(5):e1007398. Epub 2018/06/01. doi: 10.1371/journal.pgen.1007398. PubMed PMID: 29852010; PubMed Central PMCID: PMCPMC5978795.

11. Dekoninck K, Letoquart J, Laguri C, Demange P, Bevernaegie R, Simorre JP, et al. Defining the function of OmpA in the Rcs stress response. Elife. 2020;9. Epub 20200928. doi: 10.7554/eLife.60861. PubMed PMID: 32985973; PubMed Central PMCID: PMCPMC7553776.

12. Wall EA, Majdalani N, Gottesman S. IgaA negatively regulates the Rcs Phosphorelay via contact with the RcsD Phosphotransfer Protein. PLoS Genet. 2020;16(7):e1008610. Epub 2020/07/28. doi: 10.1371/journal.pgen.1008610. PubMed PMID: 32716926; PubMed Central PMCID: PMCPMC7418988.

13. Majdalani N, Heck M, Stout V, Gottesman S. Role of RcsF in signaling to the Rcs phosphorelay pathway in Escherichia coli. Journal of bacteriology. 2005;187(19):6770–8. doi: 10.1128/JB.187.19.6770-6778.2005. PubMed PMID: 16166540; PubMed Central PMCID: PMCPMC1251585.

14. Carballes F, Bertrand C, Bouche JP, Cam K. Regulation of Escherichia coli cell division genes ftsA and ftsZ by the two-component system rcsC-rcsB. Mol Microbiol. 1999;34(3):442–50. Epub 1999/11/17. PubMed PMID: 10564486.

15. Shiba Y, Miyagawa H, Nagahama H, Matsumoto K, Kondo D, Matsuoka S, et al. Exploring the relationship between lipoprotein mislocalization and activation of the Rcs signal transduction system in Escherichia coli. Microbiology (Reading). 2012;158(Pt 5):1238–48. Epub 20120209. doi: 10.1099/mic.0.056945-0. PubMed PMID: 22322964.

16. Young KD. The selective value of bacterial shape. Microbiol Mol Biol Rev. 2006;70(3):660–703. Epub 2006/09/09. doi: 70/3/660 [pii] 10.1128/MMBR.00001-06. PubMed PMID: 16959965; PubMed Central PMCID: PMC1594593.

17. Si F, Li D, Cox SE, Sauls JT, Azizi O, Sou C, et al. Invariance of Initiation Mass and Predictability of Cell Size in Escherichia coli. Curr Biol. 2017;27(9):1278–87. Epub 2017/04/19. doi: 10.1016/j.cub.2017.03.022. PubMed PMID: 28416114; PubMed Central PMCID: PMCPMC5474944.

18. Harris LK, Theriot JA. Relative Rates of Surface and Volume Synthesis Set Bacterial Cell Size. Cell. 2016;165(6):1479–92.

19. Woldringh CL, Grover NB, Rosenberger RF, Zaritsky A. Dimensional rearrangement of rod-shaped bacteria following nutritional shift-up. II. Experiments with Escherichia coli B/r. J Theor Biol. 1980;86(3):441–54. Epub 1980/10/07. PubMed PMID: 7012453.

20. Young KD. Bacterial shape: two-dimensional questions and possibilities. Annu Rev Microbiol. 2010;64:223–40. Epub 2010/09/10. doi: 10.1146/annurev.micro.112408.134102. PubMed PMID: 20825347; PubMed Central PMCID: PMCPMC3559087.

21. Taheri-Araghi S, Bradde S, Sauls JT, Hill NS, Levin PA, Paulsson J, et al. Cell-size control and homeostasis in bacteria. Current Biology. 2015;25(3):385–91.

22. Campos M, Surovtsev IV, Kato S, Paintdakhi A, Beltran B, Ebmeier SE, et al. A constant size extension drives bacterial cell size homeostasis. Cell. 2014;159(6):1433–46. Epub 2014/12/07. doi: 10.1016/j.cell.2014.11.022. PubMed PMID: 25480302; PubMed Central PMCID: PMCPMC4258233.

23. Willis L, Huang KC. Sizing up the bacterial cell cycle. Nat Rev Microbiol. 2017;15(10):606–20. Epub 2017/08/15. doi: 10.1038/nrmicro.2017.79. PubMed PMID: 28804128.

24. Holtje JV. Growth of the stress-bearing and shape-maintaining murein sacculus of Escherichia coli. Microbiol Mol Biol Rev. 1998;62(1):181–203. Epub 1998/04/08. PubMed PMID: 9529891; PubMed Central PMCID: PMC98910.

25. Gitai Z, Dye N, Shapiro L. An actin-like gene can determine cell polarity in bacteria. Proceedings of the National Academy of Sciences. 2004;101(23):8643–8.

26. Ursell T, Lee TK, Shiomi D, Shi H, Tropini C, Monds RD, et al. Rapid, precise quantification of bacterial cellular dimensions across a genomic-scale knockout library. BMC Biology. 2017;15(17):1–15.

27. Bendezu FO, de Boer PA. Conditional lethality, division defects, membrane involution, and endocytosis in mre and mrd shape mutants of Escherichia coli. J Bacteriol. 2008;190(5):1792–811. Epub 2007/11/13. doi: 10.1128/JB.01322-07. PubMed PMID: 17993535; PubMed Central PMCID: PMCPMC2258658.

28. Gitai Z, Dye NA, Reisenauer A, Wachi M, Shapiro L. MreB actin-mediated segregation of a specific region of a bacterial chromosome. Cell. 2005;120(3):329–41. Epub 2005/02/15. doi: S0092-8674(05)00045-0 [pii] 10.1016/j.cell.2005.01.007. PubMed PMID: 15707892.

29. Tropini C, Lee TK, Hsin J, Desmarais SM, Ursell T, Monds RD, et al. Principles of bacterial cell-size determination revealed by cell-wall synthesis perturbations. Cell reports. 2014;9(4):1520–7. doi: 10.1016/j.celrep.2014.10.027. PubMed PMID: 25456140; PubMed Central PMCID: PMC4254626.

30. Desmarais SM, Tropini C, Miguel A, Cava F, Monds RD, de Pedro MA, et al. High-throughput, Highly Sensitive Analyses of Bacterial Morphogenesis Using Ultra Performance Liquid Chromatography. J Biol Chem. 2015;290(52):31090–100. Epub 2015/10/16. doi: 10.1074/jbc.M115.661660. PubMed PMID: 26468288; PubMed Central PMCID: PMCPMC4692233.

31. Bi EF, Lutkenhaus J. FtsZ ring structure associated with division in Escherichia coli. 1991;354(6349):161-4. doi: 10.1038/354161a0. PubMed PMID: 1944597.

32. Dai K, Lutkenhaus J. ftsZ is an essential cell division gene in Escherichia coli. Journal of bacteriology. 1991;173(11):3500–6. PubMed PMID: 2045370; PubMed Central PMCID: PMCPMC207964.

33. Hill NS, Kadoya R, Chattoraj DK, Levin PA. Cell size and the initiation of DNA replication in bacteria. PLoS Genet. 2012;8(3):e1002549. Epub 2012/03/08. doi: 10.1371/journal.pgen.1002549. PubMed PMID: 22396664; PubMed Central PMCID: PMCPMC3291569.

34. Weart RB, Lee AH, Chien AC, Haeusser DP, Hill NS, Levin PA. A metabolic sensor governing cell size in bacteria. Cell. 2007;130(2):335–47. Epub 2007/07/31. doi: 10.1016/j.cell.2007.05.043. PubMed PMID: 17662947; PubMed Central PMCID: PMCPMC1971218.

35. Weart RB, Levin PA. Growth rate-dependent regulation of medial FtsZ ring formation. J Bacteriol. 2003;185(9):2826–34. Epub 2003/04/18. PubMed PMID: 12700262; PubMed Central PMCID: PMCPMC154409.

36. Ward Jr JE, Lutkenhaus J. Overproduction of FtsZ induces minicell formation in E. coli. Cell. 1985;42(3):941–9.

37. 37. Zietek M, Miguel A, Khusainov I, Shi H, Asmar AT, Ram S, et al. Bacterial cell widening alters periplasmic size and activates envelope stress responses. 2022.

38. Majdalani N, Hernandez D, Gottesman S. Regulation and mode of action of the second small RNA activator of RpoS translation, RprA. Mol Microbiol. 2002;46(3):813–26. Epub 2002/11/02. PubMed PMID: 12410838.

39. Clarke DJ. The Rcs phosphorelay: more than just a two-component pathway. Future Microbiol. 2010;5(8):1173–84. Epub 2010/08/21. doi: 10.2217/fmb.10.83. PubMed PMID: 20722597.

40. Jorgenson MA, Kannan S, Laubacher ME, Young KD. Dead-end intermediates in the enterobacterial common antigen pathway induce morphological defects in Escherichia coli by competing for undecaprenyl phosphate. Mol Microbiol. 2016;100(1):1–14. Epub 2015/11/26. doi: 10.1111/mmi.13284. PubMed PMID: 26593043; PubMed Central PMCID: PMCPMC4845916.

41. Moore DA, Whatley ZN, Joshi CP, Osawa M, Erickson HP. Probing for Binding Regions of the FtsZ Protein Surface through Site-Directed Insertions: Discovery of Fully Functional FtsZ-Fluorescent Proteins. J Bacteriol. 2017;199(1). Epub 2016/11/01. doi: 10.1128/JB.00553-16. PubMed PMID: 27795325; PubMed Central PMCID: PMCPMC5165096.

42. Shi H, Colavin A, Bigos M, Tropini C, Monds RD, Huang KC. Deep Phenotypic Mapping of Bacterial Cytoskeletal Mutants Reveals Physiological Robustness to Cell Size. Curr Biol. 2017;27(22):3419–29 e4. Epub 2017/11/07. doi: 10.1016/j.cub.2017.09.065. PubMed PMID: 29103935.

43. Spratt BG. Distinct penicillin binding proteins involved in the division, elongation, and shape of Escherichia coli K12. Proc Natl Acad Sci U S A. 1975;72(8):2999–3003. Epub 1975/08/01. PubMed PMID: 1103132; PubMed Central PMCID: PMCPMC432906.

44. Dominguez-Bernal G, Pucciarelli MG, Ramos-Morales F, Garcia- Quintanilla M, Cano DA, Casadesus J, et al. Repression of the RcsC-YojN-RcsB phosphorelay by the IgaA protein is a requisite for Salmonella virulence. Mol Microbiol. 2004;53(5):1437–49. Epub 2004/09/25. doi: 10.1111/j.1365-2958.2004.04213.x. PubMed PMID: 15387821.

45. Tuson HH, Auer GK, Renner LD, Hasebe M, Tropini C, Salick M, et al. Measuring the stiffness of bacterial cells from growth rates in hydrogels of tunable elasticity. Mol Microbiol. 2012;84(5):874–91. Epub 2012/05/03. doi: 10.1111/j.1365-2958.2012.08063.x. PubMed PMID: 22548341; PubMed Central PMCID: PMCPMC3359400.

46. Zheng H, Ho PY, Jiang M, Tang B, Liu W, Li D, et al. Interrogating the Escherichia coli cell cycle by cell dimension perturbations. Proc Natl Acad Sci U S A. 2016;113(52):15000–5. Epub 2016/12/14. doi: 10.1073/pnas.1617932114. PubMed PMID: 27956612; PubMed Central PMCID: PMCPMC5206551.

47. Claudi B, Sprote P, Chirkova A, Personnic N, Zankl J, Schurmann N, et al. Phenotypic variation of Salmonella in host tissues delays eradication by antimicrobial chemotherapy. Cell. 2014;158(4):722–33. doi: 10.1016/j.cell.2014.06.045. PubMed PMID: 25126781.

48. Farris C, Sanowar S, Bader MW, Pfuetzner R, Miller SI. Antimicrobial peptides activate the Rcs regulon through the outer membrane lipoprotein RcsF. J Bacteriol. 2010;192(19):4894–903. Epub 2010/08/03. doi: 10.1128/JB.00505-10. PubMed PMID: 20675476; PubMed Central PMCID: PMCPMC2944553.

49. Konovalova A, Mitchell AM, Silhavy TJ. A lipoprotein/beta-barrel complex monitors lipopolysaccharide integrity transducing information across the outer membrane. Elife. 2016;5. Epub 2016/06/11. doi: 10.7554/eLife.15276. PubMed PMID: 27282389; PubMed Central PMCID: PMCPMC4942254.

50. Gervais FG, Drapeau GR. Identification, cloning, and characterization of rcsF, a new regulator gene for exopolysaccharide synthesis that suppresses the division mutation ftsZ84 in Escherichia coli K-12. J Bacteriol. 1992;174(24):8016–22. Epub 1992/12/01. PubMed PMID: 1459951; PubMed Central PMCID: PMCPMC207539.

51. Gervais FG, Phoenix P, Drapeau GR. The rcsB gene, a positive regulator of colanic acid biosynthesis in Escherichia coli, is also an activator of ftsZ expression. J Bacteriol. 1992;174(12):3964–71. Epub 1992/06/01. PubMed PMID: 1597415; PubMed Central PMCID: PMCPMC206105.

52. Monds RD, Lee TK, Colavin A, Ursell T, Quan S, Cooper TF, et al. Systematic perturbation of cytoskeletal function reveals a linear scaling relationship between cell geometry and fitness. Cell reports. 2014;9(4):1528–37. doi: 10.1016/j.celrep.2014.10.040. PubMed PMID: 25456141.

53. Fenton AK, Gerdes K. Direct interaction of FtsZ and MreB is required for septum synthesis and cell division in Escherichia coli. EMBO J. 2013;32(13):1953–65. Epub 2013/06/13. doi: 10.1038/emboj.2013.129. PubMed PMID: 23756461; PubMed Central PMCID: PMCPMC3708099.

54. Mitosch K, Rieckh G, Bollenbach T. Noisy Response to Antibiotic Stress Predicts Subsequent Single-Cell Survival in an Acidic Environment. Cell Syst. 2017;4(4):393–403 e5. Epub 2017/03/28. doi: 10.1016/j.cels.2017.03.001. PubMed PMID: 28342718.

55. Shiver AL, Osadnik H, Peters JM, Mooney RA, Wu PI, Hu JC, et al. Chemical-genetic interrogation of RNA polymerase mutants reveals structure- function relationships and physiological tradeoffs. BioRxiv. 2020.

56. Neidhardt FC, Bloch PL, Smith DF. Culture medium for enterobacteria. J Bacteriol. 1974;119(3):736–47. Epub 1974/09/01. doi: 10.1128/JB.119.3.736-747.1974. PubMed PMID: 4604283; PubMed Central PMCID: PMCPMC245675.

57. Russell DW, Sambrook J. Molecular cloning: a laboratory manual: Cold Spring Harbor Laboratory Cold Spring Harbor, NY; 2001.

58. Edelstein AD, Tsuchida MA, Amodaj N, Pinkard H, Vale RD, Stuurman N. Advanced methods of microscope control using μManager software. Journal of biological methods. 2014;1(2):e10.

59. Schindelin J, Arganda-Carreras I, Frise E, Kaynig V, Longair M, Pietzsch T, et al. Fiji: an open-source platform for biological-image analysis. Nat Methods. 2012;9(7):676–82. Epub 2012/06/30. doi: 10.1038/nmeth.2019. PubMed PMID: 22743772; PubMed Central PMCID: PMCPMC3855844.

60. Clarke EJ, Voigt CA. Characterization of combinatorial patterns generated by multiple two-component sensors in E. coli that respond to many stimuli. Biotechnol Bioeng. 2011;108(3):666–75. Epub 2011/01/20. doi: 10.1002/bit.22966. PubMed PMID: 21246512; PubMed Central PMCID: PMCPMC3413328.

